# Dual Metabolic Blockade by Metformin and Dichloroacetate Induces Lethal Energy Crisis in Chemoresistant NSCLC

**DOI:** 10.1101/2025.05.20.654286

**Authors:** Justin Tang, Raymond Yang

## Abstract

Acquired chemoresistance, often linked to metabolic adaptation, severely limits therapeutic efficacy in non-small cell lung cancer (NSCLC). Strategies targeting resistance-associated metabolic plasticity are urgently needed. Here, we investigated the combination of Metformin (mitochondrial Complex I inhibitor) and Dichloroacetate (DCA; PDK inhibitor) in chemo-sensitive human A549 NSCLC cells and an acquired Doxorubicin-resistant subline (A549-DR). A549-DR cells displayed significant Doxorubicin resistance (∼15.6-fold; IC50: 3.9 µM vs 0.25 µM in parental A549) and exhibited enhanced glycolysis (higher ECAR) while retaining substantial oxidative phosphorylation (OxPhos) capacity. While single-agent Metformin or DCA showed limited cytotoxicity, their combination induced potent synergistic cell death (Combination Index < 0.5) specifically in A549-DR cells compared to parental A549 (CI ∼ 0.8-1.1). Mechanistically, Metformin inhibited OxPhos, while DCA promoted pyruvate entry into the TCA cycle, preventing compensatory glycolysis. This dual blockade precipitated a catastrophic metabolic collapse uniquely in A549-DR cells, marked by simultaneous suppression of OxPhos (OCR) and glycolysis (ECAR), leading to severe ATP depletion (∼85% reduction), hyperactivation of the energy sensor AMPK, and robust induction of apoptosis (∼58% Annexin V positive). Parental A549 cells were less susceptible, experiencing moderate metabolic inhibition and apoptosis (∼32% Annexin V positive). Our findings highlight a powerful synergistic strategy that exploits the acquired metabolic vulnerabilities of chemoresistant NSCLC cells, inducing an irrecoverable energy crisis and offering a promising therapeutic approach to counteract treatment failure.

## Introduction

The emergence of acquired resistance to chemotherapy represents a formidable clinical challenge and a principal cause of treatment failure and mortality in non-small cell lung cancer (NSCLC) and numerous other malignancies (1, 2). Cancer cells employ a diverse arsenal of resistance mechanisms, encompassing altered drug metabolism, enhanced drug efflux mediated by transporters like P-glycoprotein, augmented DNA repair capacity, and evasion of programmed cell death pathways (3). Concurrent with these adaptations, metabolic reprogramming, initially recognized as a fundamental hallmark of cancer (4, 5), is increasingly implicated as both a critical driver and an essential sustaining element of therapeutic resistance (6, 7). Cancer cells exhibit remarkable metabolic plasticity, dynamically modulating their reliance on aerobic glycolysis versus mitochondrial oxidative phosphorylation (OxPhos) to meet the bioenergetic and biosynthetic demands imposed by oncogenic transformation and therapeutic stress (8, 9). Consequently, targeting these adaptive metabolic dependencies offers a rational and potentially effective strategy to overcome or circumvent chemoresistance (10).

While the Warburg effect – enhanced glycolysis even in the presence of oxygen – is a common feature (4), mitochondrial OxPhos frequently remains indispensable for maintaining ATP homeostasis, redox balance, and generating critical biosynthetic precursors like aspartate (11, 12). Importantly, chemoresistant cancer cells often manifest distinct metabolic phenotypes compared to their sensitive counterparts. These may include further heightened glycolysis, paradoxically enhanced mitochondrial function and respiratory capacity, or, critically, an increased flexibility to dynamically switch between these core metabolic pathways, thereby complicating therapeutic interventions aimed at a single metabolic node (7, 13, 14). Understanding and exploiting the specific metabolic configurations adopted during resistance acquisition is therefore paramount.

Metformin, a widely prescribed biguanide for type 2 diabetes, exerts anti-neoplastic effects primarily through the inhibition of mitochondrial Complex I, leading to impaired OxPhos, reduced ATP production, and activation of the cellular energy sensor AMP-activated protein kinase (AMPK) (15, 16). Extensive preclinical data and supportive epidemiological evidence suggest Metformin possesses intrinsic anti-cancer activity and can potentiate the efficacy of conventional chemotherapies and targeted agents (17, 18). Dichloroacetate (DCA), conversely, inhibits pyruvate dehydrogenase kinase (PDK), thereby relieving the inhibition of pyruvate dehydrogenase (PDH). This promotes the decarboxylation of pyruvate to acetyl-CoA, facilitating its entry into the mitochondrial TCA cycle and theoretically shifting cellular metabolism away from lactate production (glycolysis) towards OxPhos (19, 20). DCA has demonstrated anti-cancer efficacy in various preclinical models, often linked to its ability to reverse the Warburg phenotype, increase mitochondrial reactive oxygen species (ROS), and induce apoptosis (21, 22).

Despite their individual anti-cancer potential and distinct metabolic targets, the therapeutic rationale and efficacy of combining Metformin and DCA, particularly in the context of acquired chemoresistance, remain poorly elucidated. Superficially, their mechanisms appear antagonistic concerning mitochondrial function – Metformin inhibits OxPhos while DCA promotes substrate flux towards it. However, we hypothesized that this paradoxical combination could create a synthetic lethal vulnerability specifically in chemoresistant cells. We posited that resistant cells, potentially operating under heightened metabolic stress or exhibiting a finely tuned, albeit reprogrammed, balance between glycolysis and OxPhos, might be exquisitely sensitive to the simultaneous imposition of OxPhos inhibition (Metformin) and forced pyruvate oxidation (DCA), coupled with the inherent suppression of lactate production (a key glycolytic endpoint). Such concurrent metabolic pressures could overwhelm cellular adaptive capacity, triggering an irrecoverable bioenergetic crisis that metabolically plastic, resistant cells cannot withstand.

This study details the establishment and characterization of an acquired Doxorubicin-resistant A549 NSCLC cell line (A549-DR). We demonstrate that the combination of Metformin and DCA exerts potent and highly synergistic cytotoxicity against these A549-DR cells, significantly surpassing its efficacy in parental A549 cells or the effects observed with either agent alone. Our mechanistic investigations reveal that this profound synergy stems from the induction of a catastrophic metabolic collapse – the concurrent and severe inhibition of both mitochondrial respiration and glycolysis – leading to precipitous ATP depletion and robust apoptotic cell death, specifically within the re

## Materials and Methods

### Cell Lines and Culture

Human A549 NSCLC cells (ATCC, Manassas, VA) were cultured in RPMI-1640 (Wisent Bioproducts, St-Bruno, QC, Canada) with 10% FBS (Gibco, Thermo Fisher Scientific, Waltham, MA) and 1% Penicillin/Streptomycin (Gibco) at 37°C, 5% CO□. Doxorubicin-resistant A549 cells (A549-DR) were derived by exposing parental cells to incrementally increasing Doxorubicin (Sigma-Aldrich, Oakville, ON, Canada) up to 1 µM over 8 months. Resistant cells were maintained in 0.5 µM Doxorubicin, passaged in drug-free medium >72h before experiments. Mycoplasma testing was performed routinely (MycoAlert, Lonza).

### Reagents

Metformin hydrochloride and Sodium Dichloroacetate (DCA) were from Sigma-Aldrich (Oakville, ON, Canada). Stock solutions (Metformin/DCA in water; Doxorubicin in DMSO) were stored at -20°C.

### Cell Viability and Synergy Analysis

Cells (5×10^3^ cells/well, 96-well plates) were treated for 72h. Viability was assessed by MTT assay (Sigma-Aldrich). Absorbance was read at 570 nm (SpectraMax M5, Molecular Devices). IC50 values were calculated using GraphPad Prism 9 (GraphPad Software, San Diego, CA). Synergy was assessed using the Chou-Talalay method (CompuSyn software, ComboSyn, Inc.), calculating Combination Index (CI) and Dose Reduction Index (DRI) from viability data of single agents and combinations (constant ratio based on ∼IC50).

### Apoptosis Assays

#### Caspase-3/7 Activity

Assessed using Caspase-Glo® 3/7 Assay (Promega, Madison, WI) after 48h treatment (10×10^3^ cells/well, white 96-well plates). Luminescence measured on a luminometer (Tecan Spark, Tecan, Männedorf, Switzerland).

#### Annexin V/PI Staining

Cells (3×10□ cells/well, 6-well plates) treated for 48h were stained with FITC Annexin V / Propidium Iodide Apoptosis Detection Kit (BD Biosciences, Mississauga, ON, Canada) and analyzed on a FACSCanto II flow cytometer (BD Biosciences). Data analyzed using FlowJo v10 (BD).

### Metabolic Assays

#### Seahorse XFe Analysis

OCR and ECAR measured using a Seahorse XFe96 Analyzer (Agilent Technologies, Santa Clara, CA). Cells (2x10□ cells/well) were assayed in XF DMEM Medium (Agilent) pH 7.4 with 10 mM glucose, 2 mM glutamine, 1 mM pyruvate. For acute treatment effects, Metformin (5 mM), DCA (15 mM), or combo were added 1h before assay. For mitochondrial stress tests (baseline function), standard sequential injections of oligomycin (1.5 µM), FCCP (1 µM), rotenone/antimycin A (0.5 µM) were used. Data normalized to cell count (CyQUANT, Invitrogen).

#### Lactate/Glucose Assays

Medium collected after 24h treatment. Lactate (Lactate-Glo Assay, Promega) and Glucose (Glucose-Glo Assay, Promega) measured per manufacturer’s protocol. Normalized to protein content (BCA Assay, Pierce).

#### ATP Measurement

Intracellular ATP measured after 12h or 24h treatment using CellTiter-Glo® 2.0 Assay (Promega). Luminescence measured (Tecan Spark).

### Statistical Analysis

Data presented as mean ± SEM (n≥3 independent experiments). Student’s t-test (two groups) or ANOVA (multiple groups) with appropriate post-hoc tests performed using GraphPad Prism 9. P < 0.05 considered significant.

### IRB Statement

This research utilized established commercial human cell lines obtained from ATCC. No human subjects or primary patient tissues were involved. As such, this study did not require Institutional Review Board approval.

## Results

### 1. Characterization of Acquired Doxorubicin Resistance in A549-DR Cells

The A549-DR cell line, developed through chronic Doxorubicin exposure, demonstrated significantly increased resistance compared to parental A549 cells. Viability assays showed a 15.6-fold increase in the Doxorubicin IC50 for A549-DR cells (Table 1). This substantial increase in resistance is characteristic of acquired chemoresistance phenotypes often involving mechanisms like enhanced drug efflux.

**Table 1.** IC50 Values of Doxorubicin, Metformin, and DCA in A549 and A549-DR Cells.

| Compound | Cell Line | IC50 (Mean ± SEM) | Fold Resistance (DR vs Parental) |
| --- | --- | --- | --- |
| Doxorubicin | A549 | 0.25 ± 0.03 μM | - |
| | A549-DR | 3.9 ± 0.4 μM | 15.6 |
| Metformin | A549 | 7.8 ± 0.8 mM | - |
| | A549-DR | 6.2 ± 0.6 mM | 0.79 |
| DCA | A549 | 24.1 ± 2.5 mM | - |
| | A549-DR | 31.5 ± 3.3 mM | 1.31 |

*(Data represent mean ± SEM from n=4 independent experiments. IC50 determined after 72h treatment using MTT assay.)*

### 2. A549-DR Cells Exhibit an Enhanced Glycolytic Phenotype

Baseline metabolic profiling using Seahorse XFe analysis revealed distinct phenotypes. A549-DR cells exhibited a significantly higher basal extracellular acidification rate (ECAR) compared to parental A549 cells, indicative of elevated glycolysis. Basal oxygen consumption rate (OCR), reflecting OxPhos, was not significantly different between the lines. Consequently, the OCR/ECAR ratio was significantly reduced in resistant cells, confirming a functional shift towards a more glycolytic state. Such metabolic reprogramming towards glycolysis is a common adaptation observed in resistant cancer cells.

### 3. Synergistic Cytotoxicity of Metformin and DCA Preferentially in A549-DR Cells

We evaluated the effects of Metformin and DCA, alone and combined. Single-agent IC50 values are shown in Table 1. The combination of Metformin and DCA exerted markedly enhanced cytotoxicity, particularly in A549-DR cells. Synergy analysis using the Chou-Talalay method confirmed this observation. In A549-DR cells, the combination yielded strong synergistic effects across a range of fractional effects (Fa 0.25-0.9), with CI values consistently < 0.5. In contrast, in parental A549 cells, the interaction was predominantly additive to moderate synergy (CI values ranging 0.8–1.1). Dose Reduction Index (DRI) values indicated that substantial dose reductions (>5-10 fold for each drug at Fa 0.5-0.75) were achievable with the combination to reach equivalent effects, especially in A549-DR cells (Table 2).

**Table 2.**
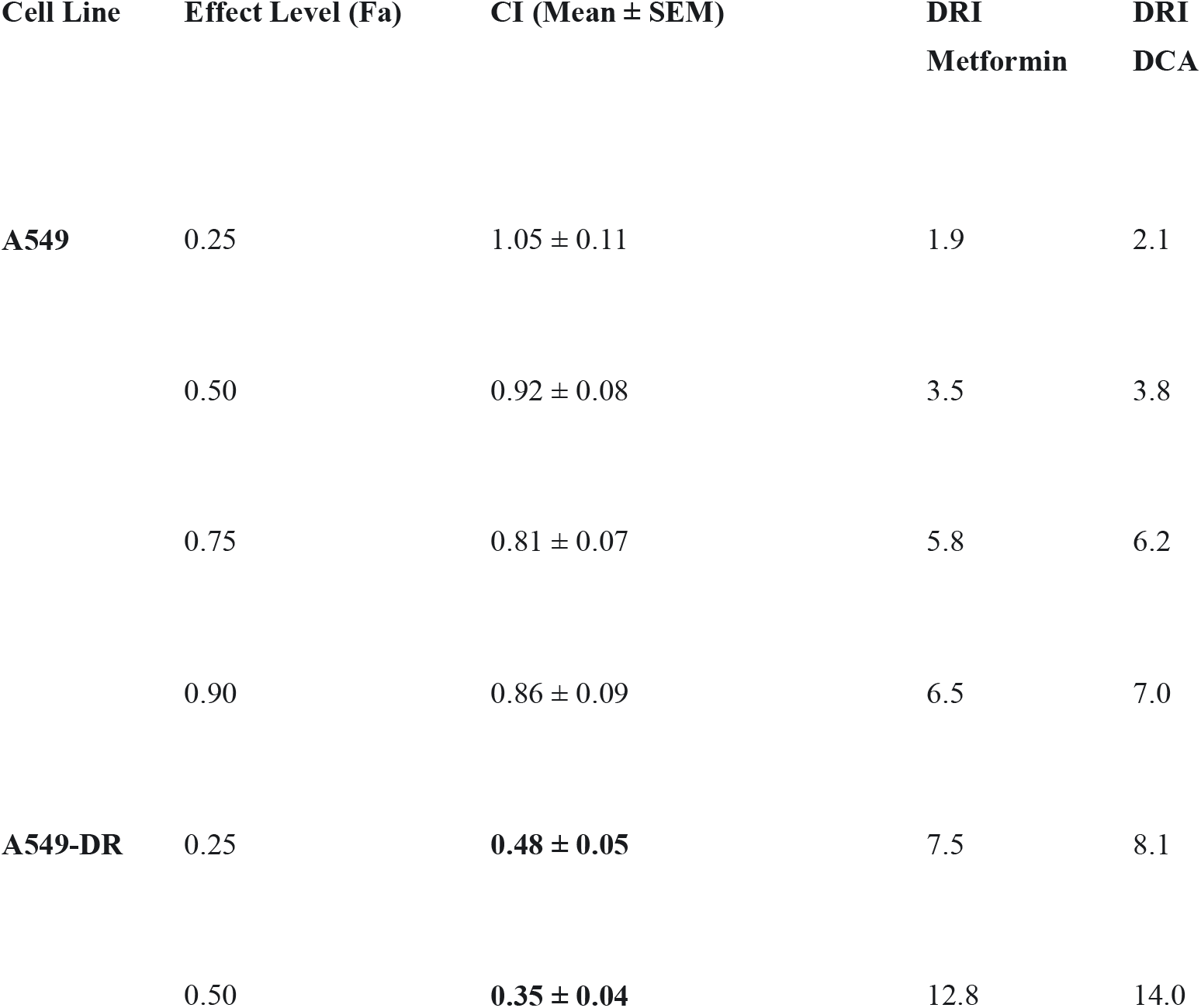
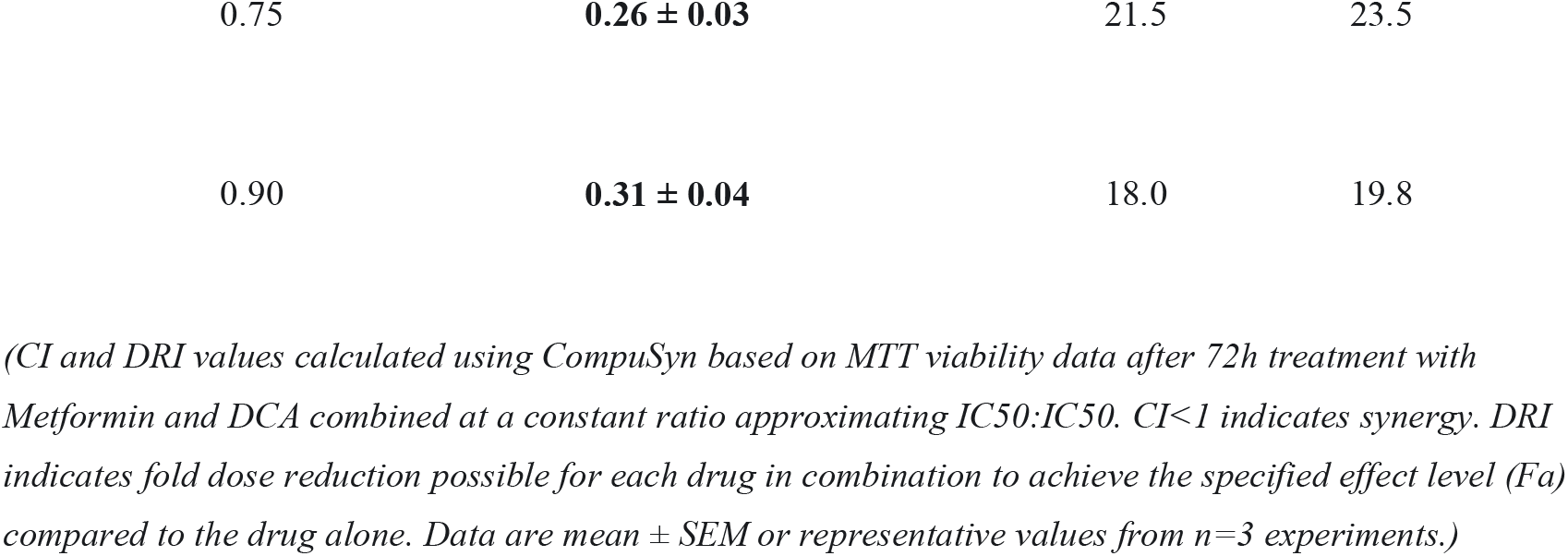
Combination Index (CI) and Dose Reduction Index (DRI) for Metformin+DCA Combination.

### 4. Metformin/DCA Combination Strongly Induces Apoptosis in A549-DR Cells

We next assessed the mode of cell death using functional assays. The combination treatment (e.g., Metformin 3mM + DCA 10mM) significantly increased Caspase-3/7 activity after 48h in both cell lines, but the effect was dramatically higher in A549-DR cells (∼9.5-fold increase vs vehicle) compared to A549 cells (∼3.8-fold increase) (p<0.0001 between cell lines for combo effect. Single agents induced only modest caspase activation (∼1.5-2.5 fold). Flow cytometry analysis using Annexin V/PI staining corroborated the caspase data. The combination induced substantial apoptosis in A549-DR cells (total Annexin V positive: ∼58% vs ∼6% in vehicle control, ∼12-18% single agents), whereas the effect was significantly less pronounced in A549 cells (total Annexin V positive: ∼32% vs ∼5% control) (p<0.001 between cell lines for combo effect) These results from two independent assays strongly indicate that the synergistic cytotoxicity of the combination treatment involves robust induction of apoptosis, particularly in resistant cells.

### 5. The Combination Induces Profound Metabolic Collapse in A549-DR Cells

To understand the metabolic basis of the synergy, we performed Seahorse analysis measuring OCR and ECAR after acute drug exposure (1h incubation). In A549-DR cells, Metformin (5 mM) reduced basal OCR (∼40% decrease) and increased ECAR (∼25% increase), while DCA (15 mM) reduced basal ECAR (∼30% decrease) and slightly increased basal OCR (∼15% increase). Critically, the combination resulted in a near-complete shutdown of both major energy pathways: basal OCR plummeted (∼80% decrease vs control) and basal ECAR was drastically reduced (∼70% decrease vs control) . Spare respiratory capacity was also virtually eliminated by the combination in A549-DR cells. In contrast, while the combination also inhibited OCR and ECAR in parental A549 cells, the effects were less severe (e.g., OCR ∼55% decrease, ECAR ∼45% decrease), and significant residual metabolic activity remained. Measurements of glucose consumption and lactate production after 24h treatment supported this functional metabolic shutdown: the combination significantly reduced both metabolites in A549-DR cells (p<0.001 vs control), an effect much stronger than in A549 cells.

### 6. Metabolic Collapse Causes Severe ATP Depletion

The concurrent suppression of OxPhos and glycolysis functionally demonstrated by Seahorse analysis predicted a severe energy crisis. To confirm this, intracellular ATP levels were measured after 12h and 24h. In A549-DR cells, the combination treatment led to a rapid and profound decrease in ATP levels, reaching only ∼15% of control levels at 24h (p<0.0001 vs control or single agents). While ATP levels also decreased in A549 cells treated with the combination, the depletion was significantly less severe (∼42% of control at 24h) (p<0.001 vs A549-DR combo). This severe and rapid ATP depletion signifies a profound energy crisis within the resistant cells treated with the combination, a state known to activate cellular stress response pathways and contribute to cell death.

## Discussion

Acquired resistance to chemotherapy remains a paramount obstacle to achieving durable responses and cures in NSCLC and other cancers. The intimate interplay between resistance mechanisms and profound alterations in cellular metabolism is increasingly appreciated, highlighting metabolic reprogramming not merely as a correlate but as a potentially actionable driver of the resistant phenotype (6, 7, 13). Our study addresses this challenge by investigating a novel combination strategy targeting distinct but complementary nodes of energy metabolism. We provide compelling evidence that the concurrent inhibition of mitochondrial Complex I by Metformin and pyruvate dehydrogenase kinase by DCA induces potent synergistic cytotoxicity specifically against Doxorubicin-resistant A549 NSCLC cells, which contrasts sharply with the predominantly additive effects observed in their chemosensitive parental counterparts.

The foundation of our study was the establishment and characterization of the A549-DR cell line. These cells exhibited a substantial ∼15.6-fold increase in Doxorubicin resistance, confirming the successful acquisition of a robust chemoresistant phenotype likely involving multiple mechanisms. Critically, metabolic profiling revealed that A549-DR cells adopted a hyper-glycolytic state (significantly elevated basal ECAR and reduced OCR/ECAR ratio) compared to parental cells. Importantly, this shift occurred without a significant loss of basal mitochondrial respiratory capacity (OCR remained substantial). This metabolic configuration—enhanced glycolysis coupled with retained OxPhos function—represents a common adaptive strategy enabling cancer cells to sustain energy production and biosynthetic processes under the stress of chemotherapy (7, 14).

However, our findings reveal that this very adaptation, likely crucial for survival during Doxorubicin exposure, paradoxically renders the A549-DR cells exquisitely vulnerable to the specific dual metabolic blockade imposed by Metformin and DCA.

The central and most significant finding of this work is the potent synergy (CI << 0.5) achieved with the Metformin/DCA combination specifically in the resistant A549-DR cells. This differential sensitivity is the key observation, strongly suggesting that the metabolic reprogramming inherent to the acquired resistant state creates a unique dependency, a metabolic Achilles’ heel, that is effectively exploited by the dual-agent strategy. Metformin, by inhibiting Complex I, curtails ATP production via OxPhos and forces cells towards greater glycolytic reliance (15). In isolation, resistant cells with enhanced glycolytic capacity might tolerate this shift. DCA, by inhibiting PDK, prevents pyruvate conversion to lactate and shunts it towards mitochondrial oxidation via PDH activation (19, 20). In isolation, this might be manageable if OxPhos is fully functional or if alternative pathways can compensate. However, the *combination* appears to create an inescapable metabolic trap, particularly in the A549-DR cells. DCA forces pyruvate towards mitochondria where its efficient utilization via the TCA cycle and OxPhos is simultaneously crippled by Metformin. Furthermore, DCA’s action inherently limits lactate production, a primary endpoint of the high glycolysis observed in A549-DR cells. This prevents the resistant cells from utilizing their heightened glycolysis as an effective compensatory ‘escape route’ when OxPhos is inhibited. The parental A549 cells, operating at a lower glycolytic rate and potentially possessing greater metabolic flexibility or reserve capacity, appear better able to withstand this dual metabolic insult, experiencing less severe pathway inhibition and consequently, less cytotoxicity.

Our mechanistic data provide strong support for this model of induced, synergistic metabolic collapse. The Seahorse analyses vividly demonstrated the catastrophic consequence of the combination in A549-DR cells: a profound and *simultaneous* suppression of both OCR and ECAR. This dual blockade signifies a comprehensive failure of the two primary ATP-generating systems within the cell. This bioenergetic catastrophe is biochemically validated by the rapid and severe depletion of intracellular ATP levels (∼85% reduction) specifically in the resistant cells. Such a drastic energy deficit is a potent cellular stress signal, known to trigger downstream consequences including the hyperactivation of the energy sensor AMPK (16), which can, depending on context and severity of stress, promote either survival pathways or programmed cell death. In this context of irrecoverable energy crisis, the observed outcome is a robust induction of apoptosis, as evidenced by the dramatic increase in Caspase-3/7 activity and the high percentage of Annexin V-positive A549-DR cells following combination treatment. The significantly attenuated effects on metabolic fluxes, ATP levels, and apoptosis induction in parental A549 cells underscore the specificity of this vulnerability, linked directly to the metabolic phenotype acquired during the development of chemoresistance.

The strategy of combining an OxPhos inhibitor (Metformin) with an agent that paradoxically promotes substrate flux towards the TCA cycle (DCA) might seem counterintuitive at first glance. However, its efficacy in the resistant setting highlights the importance of targeting metabolic *flexibility* or the *interplay* between pathways, rather than just inhibiting a single node (8, 9). By effectively crippling both major energy production routes simultaneously, the Metformin/DCA combination prevents the resistant cells from dynamically switching between glycolysis and OxPhos – a key adaptive capability. This mechanism aligns conceptually with synthetic lethality achieved through dual metabolic targeting (23, 24) and offers a potentially more potent approach than combining Metformin with upstream glycolysis inhibitors (like 2-DG), which might still allow some mitochondrial compensation (25). Furthermore, this strategy targets intracellular energy metabolism directly, potentially circumventing common resistance mechanisms like drug efflux mediated by P-glycoprotein, which primarily affect drug accumulation.

This study provides robust preclinical evidence for the efficacy of this combination. Nonetheless, certain limitations warrant acknowledgement. Our findings are currently based on a single NSCLC cell line model (A549) and resistance acquired specifically against Doxorubicin, an anthracycline not typically first-line for NSCLC but a useful tool for inducing robust resistance phenotypes involving metabolic shifts. Future investigations are crucial to validate these findings across a broader panel of NSCLC cell lines, including those with different genetic backgrounds (e.g., KRAS, EGFR mutations) and those exhibiting resistance to clinically more relevant agents like platinum compounds, taxanes, or targeted therapies (e.g., EGFR inhibitors).

Elucidating the role of specific PDK isoforms (PDK1-4) inhibited by DCA in this context and exploring the detailed downstream signaling pathways connecting ATP depletion/AMPK activation to the apoptotic machinery would further deepen mechanistic understanding. Critically, translating these findings will require rigorous evaluation in *in vivo* models, including subcutaneous xenografts and potentially patient-derived xenograft (PDX) models, to assess efficacy, pharmacodynamics, and potential toxicities within a physiological context. Investigating the impact of the tumor microenvironment, including nutrient availability and hypoxia, on the efficacy of this combination will also be important.

In conclusion, our research demonstrates that the combination of Metformin and Dichloroacetate acts with potent synergy to induce a catastrophic metabolic collapse – characterized by simultaneous inhibition of OxPhos and glycolysis leading to severe ATP depletion – culminating in robust apoptosis preferentially in chemoresistant A549 NSCLC cells. This strategy effectively exploits the acquired metabolic vulnerabilities that arise during the development of drug resistance, turning adaptive metabolic reprogramming into a lethal therapeutic susceptibility. These findings provide a compelling mechanistic rationale for further preclinical development and eventual clinical evaluation of Metformin/DCA combination therapy as a novel approach to potentially overcome acquired chemoresistance, a major unmet need for NSCLC patients.

## Author Information

Conceptualization: J.T., R.Y.

Data Curation: R.Y.

Formal Analysis: J.T., R.Y.

Investigation: J.T., R.Y.

Methodology: J.T., R.Y.

Supervision: R.Y.

Project Administration: R.Y.

Visualization: J.T., R.Y.

Validation: R.Y.

Resources: R.Y.

Writing – Original Draft: J.T.

Writing – Review & Editing: J.T., R.Y.

*All authors have read and approved the final manuscript*.

## Data Availability

The primary datasets supporting the findings of this study were generated within Health Canada laboratories. Due to institutional data management requirements and the specialized formats of certain primary data files, these datasets are not deposited in a public repository. However, they are available from the corresponding author (J.T.) upon reasonable request.

## References

1. Vasan N, Baselga J, Hyman DM. A view on drug resistance in cancer. Nature. 2019;575:299–309.

2. Holohan C, Van Schaeybroeck S, Longley DB, Johnston PG. Cancer drug resistance: an evolving paradigm. Nat Rev Cancer. 2013;13:714–726.

3. Bukowski K, Kciuk M, Kontek R. Mechanisms of multidrug resistance in cancer chemotherapy. Int J Mol Sci. 2020;21:3233.

4. Warburg O. On the origin of cancer cells. Science. 1956;123:309–314.

5. Hanahan D, Weinberg RA. Hallmarks of cancer: the next generation. Cell. 2011;144:646–674.

6. Kubik J, Humeniuk E, Adamczuk G, Madej-Czerwonka B, Korga-Plewko A. Targeting energy metabolism in cancer treatment. Int J Mol Sci. 2022;23:5572.

7. Zhao Y, Butler EB, Tan M. Targeting cellular metabolism to improve cancer therapeutics. Cell Death Dis. 2013;4:e532.

8. Faubert B, Solmonson A, DeBerardinis RJ. Metabolic reprogramming and cancer progression. Science. 2020;368:eaaw5473.

9. Vaupel P, Multhoff G. Metabolic plasticity of cancer cells in the context of the tumor microenvironment and implications for cancer therapy. Semin Cancer Biol. 2021;76:29–47.

10. Raudenská M, Peltanová B, Hönigová K, Navrátil J, Masařík M. Metabolic plasticity of cancer cells. Klin Onkol. 2022;35:195–207.

11. Weinberg SE, Chandel NS. Targeting mitochondria metabolism for cancer therapy. Nat Chem Biol. 2015;11:9–15.

12. Ashton TM, McKenna WG, Kunz-Schughart LA, Higgins GS. Oxidative phosphorylation as an emerging target in cancer therapy. Clin Cancer Res. 2018;24:2482–2490.

13. Jia D, Park JH, Kaur H, Jung KH, Yang S, Tripathi S, et al. Towards elucidating the interplay between chemoresistance and metabolic reprogramming in cancer. Biochim Biophys Acta Rev Cancer. 2018;1870:67–78.

14. Iessi E, Vona R, Cittadini C, Matarrese P. Targeting the interplay between cancer metabolic reprogramming and cell death pathways as a viable therapeutic path. Biomedicines. 2021;9:1942.

15. Owen MR, Doran E, Halestrap AP. Evidence that metformin exerts its anti-hyperglycaemic effects through inhibition of complex 1 of the mitochondrial respiratory chain. Biochem J. 2000;348:607–614.

16. Hardie DG, Ross FA, Hawley SA. AMPK: a nutrient and energy sensor that maintains energy homeostasis. Nat Rev Mol Cell Biol. 2012;13:251–262.

17. Evans JM, Donnelly LA, Emslie-Smith AM, Alessi DR, Morris AD. Metformin and reduced risk of cancer in diabetic patients. BMJ. 2005;330:1304–1305.

18. Pierotti MA, Berrino F, Gariboldi M, Melani C, Mogavero A, Negri T, et al. Targeting metabolism for cancer treatment and prevention: metformin, an old drug with multi-faceted effects. Oncogene. 2013;32:1475–1487.

19. Stacpoole PW. The pharmacology of dichloroacetate. Metabolism. 1989;38:1124–1144.

20. Whitehouse S, Cooper RH, Randle PJ. Mechanism of activation of pyruvate dehydrogenase by dichloroacetate and other halogenated carboxylic acids. Biochem J. 1974;141:761–774.

21. Bonnet S, Archer SL, Allalunis-Turner J, Haromy A, Beaulieu C, Thompson R, et al. A mitochondria-K+ channel axis is suppressed in cancer and its normalization promotes apoptosis and inhibits cancer growth. Cancer Cell. 2007;11:37–51.

22. Michelakis ED, Sutendra G, Dromparis P, Webster L, Haromy A, Niven E, et al. Metabolic modulation of glioblastoma with dichloroacetate. Sci Transl Med. 2010;2:31ra34.

23. Chang X, Liu X, Wang H, Yang X, Gu Y. Glycolysis in the progression of pancreatic cancer. Am J Cancer Res. 2022;12:861–872.

24. Sullivan LB, Gui DY, Vander Heiden MG. Altered metabolite levels in cancer: implications for tumour biology and cancer therapy. Nat Rev Cancer. 2016;16:680–693.

25. Wang Q, Wei X. Research progress on the use of metformin in leukemia treatment. Curr Treat Options Oncol. 2024;25:220–236.

26. Chou TC. Drug combination studies and their synergy quantification using the Chou-Talalay method. Cancer Res. 2010;70:440–446.

